# Frontal mechanisms underlying primate calls recognition by humans

**DOI:** 10.1101/2023.02.13.528425

**Authors:** Leonardo Ceravolo, Coralie Debracque, Eva Pool, Thibaud Gruber, Didier Grandjean

## Abstract

The ability to process verbal language seems unique to humans and relies not only on semantics but on other forms of communication such as affective vocalisations, that we share with other primate species—particularly great apes (*Hominidae*). To better understand these processes at the behavioural and brain level, we asked human participants to categorize vocalizations of four primate species including human, great apes (chimpanzee and bonobo), and monkey (rhesus macaque) during MRI acquisition. Classification was above chance level for all species but bonobo vocalizations. Imaging analyses were computed using a participant-specific, trial-by-trial fitted probability categorization value in a model-based style of data analysis. Model-based analyses revealed the implication of the bilateral orbitofrontal cortex and inferior frontal gyrus *pars triangularis* (IFG_tri_) respectively correlating and anti-correlating with the fitted probability of accurate species classification. Further conjunction analyses revealed enhanced activity in a sub-area of the left IFG_tri_ specifically for the accurate classification of chimpanzee calls compared to human voices. Our data therefore reveal distinct frontal mechanisms that shed light on how the human brain evolved to process non-verbal language.

**Author contributions:** CD and LC were involved in every steps of the study including experimental design, programming, data acquisition, data analysis and redaction of the first draft of the manuscript and subsequent editing. EP helped design the model-based MRI analyses and interpret the results. TG and DG were involved in the experimental design and study planification. All authors wrote and edited the manuscript.

## Introduction

Processing and understanding non-verbal language are fundamental aspects of everyday life in human (1) and animal (2, 3) communication. Their importance can sometimes even be greater than that of verbal communication since a short, powerful in-context vocal burst, for instance, an urgency signal, can easily be noticed and understood by other individuals (2). A critical part of non-verbal language can therefore be represented by ‘affective’ communication, especially through affective vocalizations, which are of particular interest for cross-species research. Indeed, the ability to express and understand such vocalizations is shared by several primate species, particularly Hominids, a taxonomic family that includes humans and other great apes (4, 5). Such commonalities therefore open a critical window into the investigation of human brain structures recruited by non-verbal communication, for instance by studying these crucial abilities to perceive, process and subsequently categorize and classify affective vocalizations expressed by primate species. These abilities are known to recruit superior temporal, orbitofrontal and inferior frontal cortices in humans (6–9) while these brain areas evolved a lot in primates (10, 11). We will now detail the particular importance of studying cross-species non-verbal language processing and categorization taking place in the frontal cortex of humans.

The *Hominidae* clade appeared between 13 and 18 million years ago (12). Encompassing all living great apes including humans (*Homo sapiens*), chimpanzees (*Pan troglodytes*), bonobos (*Pan Paniscus*), gorillas (*Gorilla subs*), and orangutans (*Pongo subs*), as well as our extinct ancestors, this unique primate taxon is key to understanding the evolution of human behaviour, physiology, communication and cognitive abilities. Very few studies have used bonobo vocalizations as stimuli (13), despite sharing the same phylogenetic proximity to *Homo sapiens* with chimpanzees, their behaviour repertoire—of which gestures can be decoded by humans (14)—including their vocal communication are noticeably different (15–18). Despite a larger and more folded brain compared to other great ape species, human neuroanatomical traits are mainly considered to belong to a continuum in the primate brain evolution (11, 19). For instance, findings in anatomical magnetic resonance imaging (MRI) have demonstrated the existence of a large frontal cortex in all great ape species— humans included(11)—emphasizing the particular interest of a comparative approach to investigate anatomical structures and related functions of the frontal regions. Moreover, the functions of the frontal lobe, often associated to problem solving, emotion processing and especially evaluative judgment, communication and language or even motor and sexual behaviours (6, 20–25), as well as their related brain structures are shared by most primate species. This includes two critical brain structures that are the focus of this article: the orbitofrontal cortex (OFC) and the inferior frontal gyrus (IFG).

Existing in phylogenetically distant non-human primate species such as rhesus macaques (*Macaca mulatta*) (26), the OFC has similar neuroanatomical architecture and functions across the primate family tree. From humans to monkeys, studies have highlighted the roles of OFC in emotional memory (20, 21), emotional expression (9, 27), executive functions (20) and integration of contextual information in emotional evaluation (8). Interestingly, despite these shared evolutionary OFC functions in primate species, its role in heterospecific affective recognition, such as affective vocalizations, has rarely been investigated. In fact, recent functional MRI data have pointed out a greater enhancement of activity in OFC, in particular of its medial part, in the human identification of emotional voices compared to affective animal vocalizations including positive and negative calls expressed by chimpanzees, macaques and cats (*Felis catus*) (28, 29). More investigations on the role of the human OFC in this context are thus crucially needed.

Functionally and anatomically connected to the OFC (30), the IFG is also present in great apes as well as in monkey species (31). Yet, unlike the OFC, the left IFG is most prominently linked to speech production (Broca’s area) in humans (32), the only species ‘capable’ of a complex and sophisticated language. Hence, in addition to likely shared functions such as decision formation (6, 33), inhibition (34, 35) and emotional judgement (8, 36–38) shaped by natural selection, the IFG also became specialized in semantic auditory processing in the human lineage in the course of evolution (7, 39, 40). Because the current roles of human IFG are linked to both the *Hominidae* and the specific *Homo sapiens* evolutionary histories (41), this frontal region, together with the OFC, is of high interest to investigate conspecific and heterospecific identification mechanisms. Noteworthy is the fact that these specific frontal regions have strong anatomical ties with temporal lobe regions that are crucial in processing vocalizations, especially when they are biologically relevant (23, 30, 36, 42). Besides the importance of OFC and IFG regions in the evolution of primate brains, comparative and integrative approaches to investigate the current functions of these specific frontal regions are still missing. This is especially true for auditory affective processing (43), species categorization and by extension non-verbal communication comprehension in humans—excluding sign language.

The present study focused on the processing and subsequent categorization of primate vocalizations in human OFC and IFG using fMRI. To do so, we asked participants to categorize four primate species (human, chimpanzee, bonobo, and macaque) through their affective vocalizations. Non-verbal sounds—namely affective vocal bursts—were selected as human vocalizations to match as well as possible the nature of our primates’ vocalizations stimuli. Research on vocal production and perception has indeed demonstrated that non-verbal vocalizations expressed by humans are similar to those expressed by other mammalian species, non-human primates included, especially in affiliative (e.g., play) or agonistic contexts (e.g., aggression) (44). Interestingly, as most animal vocalizations are expressed in a motivational or affective context, existing literature has remained focused on the human recognition of affects in heterospecific vocalizations while neglecting humans’ ability to identify other species independently of affect (13, 29, 45–49). Hence, in light of previous literature as well as taking into account the commonalities and differences of the selected four primate species in their evolutionary path, we predicted: i) a behavioural gradient reflecting phylogenetic distance from humans for the species with a higher probability of correctly categorizing human, then chimpanzee, bonobo and finally macaque affective vocalizations (13, 46, 48); for similar reasons, we predicted that chimpanzee and bonobo vocalizations should be more confused with each other than with macaque calls; ii) neural correlates of the probability of correctly categorizing the four primate species in the OFC and IFG, especially in the medial OFC (28, 29) and IFG_tri/op_(_50_), respectively; iii) a gradient of activity in the *pars triangularis* and *opercularis* of the IFG when comparing the processing and correct categorization of each non-human primate species’ vocalizations to human voices using conjunction analyses, both subparts being crucially involved in this process (37, 50).

## Results

The present study aimed at assessing the ability of humans to accurately classify human and more importantly other primate calls, in the present case chimpanzee, bonobo, and macaque vocalizations (Fig.1**a**) and how frontal brains areas may underlie such successful non-verbal stimuli categorization. Behavioural analyses were performed using a Species factor (human, chimpanzee, bonobo, macaque) interacting with the Affective context factor for each vocalization (affiliative, agonistic ‘threat’, agonistic ‘distress’) in a generalized linear mixed-effects modelling of the accuracy data (‘1’, correct response; ‘0’, incorrect response) and a linear mixed-effects modelling of the confusion data. This statistical procedure was used to better assess the probability of correctly categorizing each species’ vocalizations as well as the pattern of confusion between each species, respectively. Following similar reasoning, we used the trial-level probability of correct species categorization in model-based fMRI methodology. We tested the specific frontal mechanism underlying the combined processing and accurate classification of each non-human primate species vocalizations compared to human voices using conjunction analyses. Both analyses were conducted in our regions of interest (ROI), namely in the bilateral IFG (*pars opercularis*, *triangularis*, *orbitalis*) and OFC (superior, middle, medial) using masked second-level analyses with voxelwise corrected statistics (*p*<.05 False Discovery Rate correction ‘FDR’, k>5 voxels). Selected low-level acoustics (mean and standard deviation of stimulus energy and fundamental frequency) were used as covariates of no interest for behavioural and neuroimaging analyses to better control our data for low-level vocal fluctuations. Statistical modelling power is reported for fixed effects (‘R^2^c’) and for both fixed and random effects taken together (‘R^2^m’).

## Behavioural results

### Accuracy data

As predicted, the generalized linear mixed-effects on accuracy revealed a significant effect of Species factor (χ^2^(3)=222.78, *p*<.0001, R^2^c=56.38% and R^2^m=61.10%). All complementary results of this model can be found in Supplementary Table 1. We then tested the contrasts of interest according to our behavioural hypothesis namely the impact of phylogenetic proximity on accurate species categorization: our results highlighted a higher probability of correctly categorizing human compared to the vocalizations of each non-human primate (χ^2^(1)=128.96, *p_Bonf_* <.0001; human > chimpanzee: χ (1)=95.89, *p_Bonf_* <.0001; human > bonobo: χ (1)=199.51, *p_Bonf_* <.0001; human > macaque: χ (1)=79.40, *p_Bonf_* <.0001). We additionally compared each non-human primate species with each other and found a higher probability of correct categorization for chimpanzee compared to bonobo (χ^2^(1)=52.64, *p_Bonf_* <.0001), and macaque compared to bonobo vocalizations (χ^2^(1)=18.99, *p_Bonf_* <.0001). No difference was observed when comparing the probability of correct species categorization between chimpanzee and macaque vocalizations (χ^2^(1)=1.84, *p_Bonf_* =.17, Fig. 1b).

**Figure 1.**
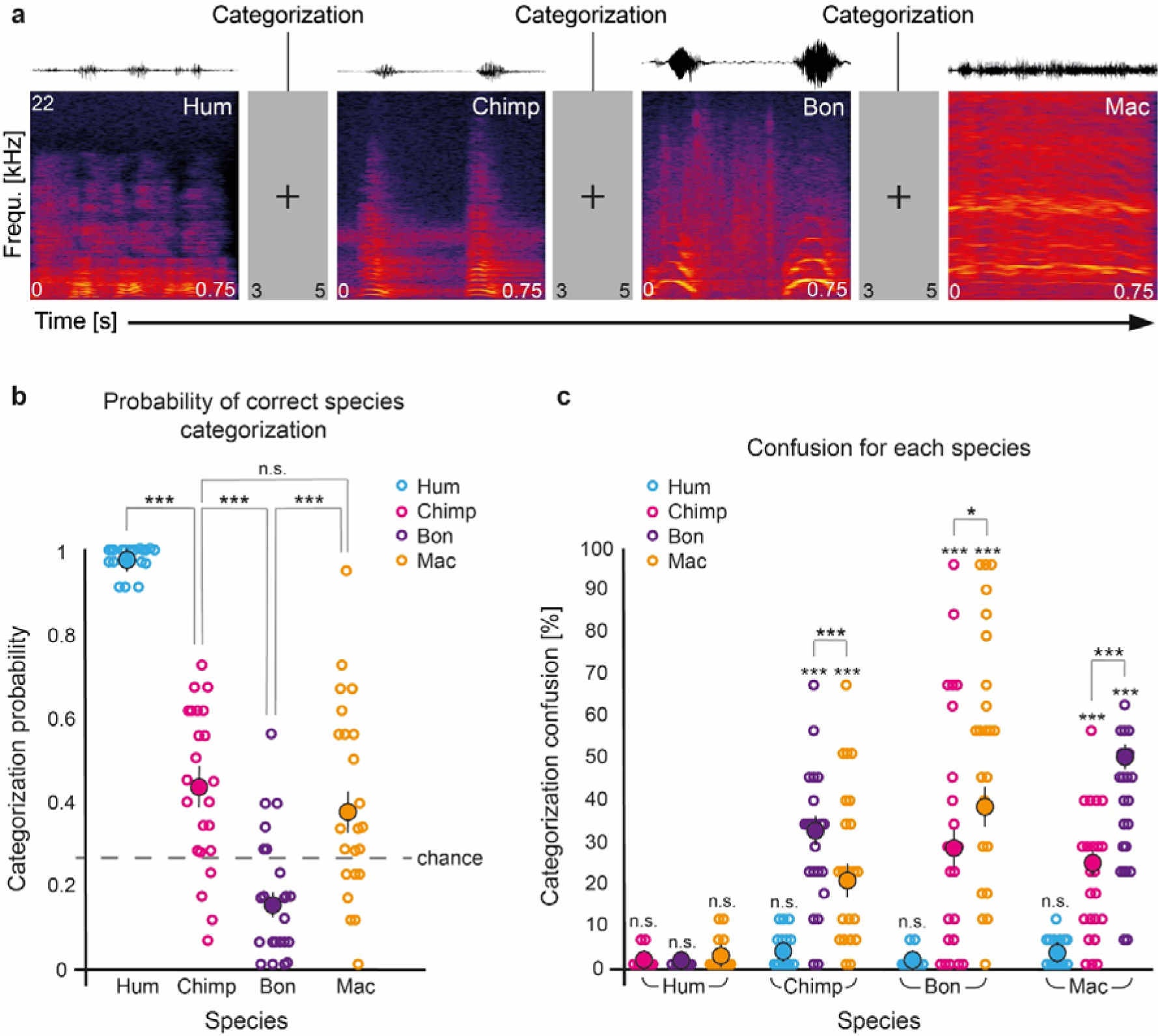
Experimental procedure and behavioural results of the probability of correct species categorization and confusions. **a,** Example of four consecutive experimental task trials (inter-trial interval of 3-5s) for which the participants had to categorize the species’ stimulus (750ms duration) for each trial (pseudo-randomized species order) and spectrogram example for each species. **b,** Probability of the correct categorization of each species (*y* axis) modelled across all participants, trials and species (i.e., human, chimpanzee, bonobo, macaque); Each circle represents a participant’s averaged value per species, full circles represent the general average per species across participants ± the standard error of the mean, chance level is 25% (grey dotted line). **c,** Percentage of categorization confusion (*y* axis) specific to each species (*x* axis); Each circle represents a participant’s averaged value, full circles represent the general average per species across participants ± the standard error of the mean. Hum: human; Chimp: chimpanzee; Bon: bonobo; Mac: macaque. \*\*\**p*<.001, \**p*<.05, n.s.: non-significant.

Confusion data analyses were computed by considering species miscategorization, namely one species’ vocalization being categorized as belonging to another species (Fig.1**c**). The confusion for human voices was not significant (χ^2^(3)=1.65, *p*=.65), meaning that these were not significantly miscategorised with any other species. Confusion concerning chimpanzee vocalizations showed a significant main effect of Species (χ^2^(3)=265.79, *p*<.0001; R^2^c=16.41% and R^2^m=24.64%), showing that these were mostly miscategorised as bonobo vocalizations (χ^2^(1)=149.52, *p_Bonf_* <.0001), and then as macaque (χ^2^(1)=47.11, *p_Bonf_* <.0001) vocalizations (for chimpanzee confusions, bonobo > macaque: χ^2^(1)=15.34, *p_Bonf_* <.0001), but not as human voices (χ^2^(1)=0.21, *p_Bonf_*>.99). Confusion for bonobo vocalizations showed a significant main effect of the Species factor (χ^2^(3)=107.96, *p*<.0001; R^2^c=24.50% and R^2^m=28.95%) and specific confusion was the highest with macaque (χ^2^(1)=114.70, *p_Bonf_*<.0001) and then chimpanzee (χ^2^(1)=111.28, *p_Bonf_* <.0001) vocalizations (for bonobo confusions, macaque > chimpanzee: χ^2^(1)=5.33, *p_Bonf_* <.05) but no significant confusion with human voices was observed (χ^2^(1)=0.06, *p_Bonf_* >.99). Confusion for macaque vocalizations showed a main effect of the Species factor (χ^2^(3)=469.68, *p*<.0001; R^2^c=29.01 % and R^2^m=30.65%) with highest confusion with bonobo (χ^2^(1)=565.04, *p_Bonf_* <.0001) followed by chimpanzee (χ^2^(1)=145.96, *p_Bonf_* <.0001) vocalizations (for macaque confusions, bonobo > chimpanzee: χ^2^(1)=87.34, *p_Bonf_* <.0001). No confusion with the human voice reached significance (χ^2^(1)=0.04, *p_Bonf_* >.99). All other specific confusion effects can be found in Supplementary Table 2.

### Neuroimaging results using IFG and OFC as regions of interest

We first used a model-based approach in order to characterize the general frontal mechanism underlying the accurate categorization of primate vocalizations by human participants (see model 1 in the methods section). To do so, we modelled all trials with the sample-specific fitted probability of a correct vs. incorrect species categorization as a covariate of interest for each trial’s onset. This covariate was then taken as the main regressor of interest to masked group-level analyses to reveal frontal patterns of activity related to the general human neural mechanism of classifying human, chimpanzee, bonobo, and macaque vocalizations. These fitted probability values were obtained across participants and species by predicting accuracy data at the trial level, therefore including all trials, correct and incorrect, in a logistic regression analysis. Fixed effects included the Species and Affective context factors and covariates of no interest (mean and standard deviation of vocalization energy and fundamental frequency) with random effects characterizing the identity of each participant (random intercept) and each stimulus (random slope). Activity correlating with the probability of correctly categorizing each species’ vocalizations was revealed in the right IFG_tri_ (Fig.2**b**) and bilateral IFG_orb_ (Fig.2**abd**) as well as in the bilateral OFC_med_ (Fig.2**c**). Anti-correlated activity was revealed in a large left IFG_tri_, IFG_op_ cluster (Fig.2**ef**) and to a lesser extent in the left IFG_orb_ (Fig.2**e**), right IFG_op_ and IFG_orb_ (Fig.2**fh**) and in the right OFC_mid_ (Fig.2**b**) and left OFC_sup_ (Fig.2**e**). See Supplementary Table 3 for cluster coordinates.

**Figure 2.**
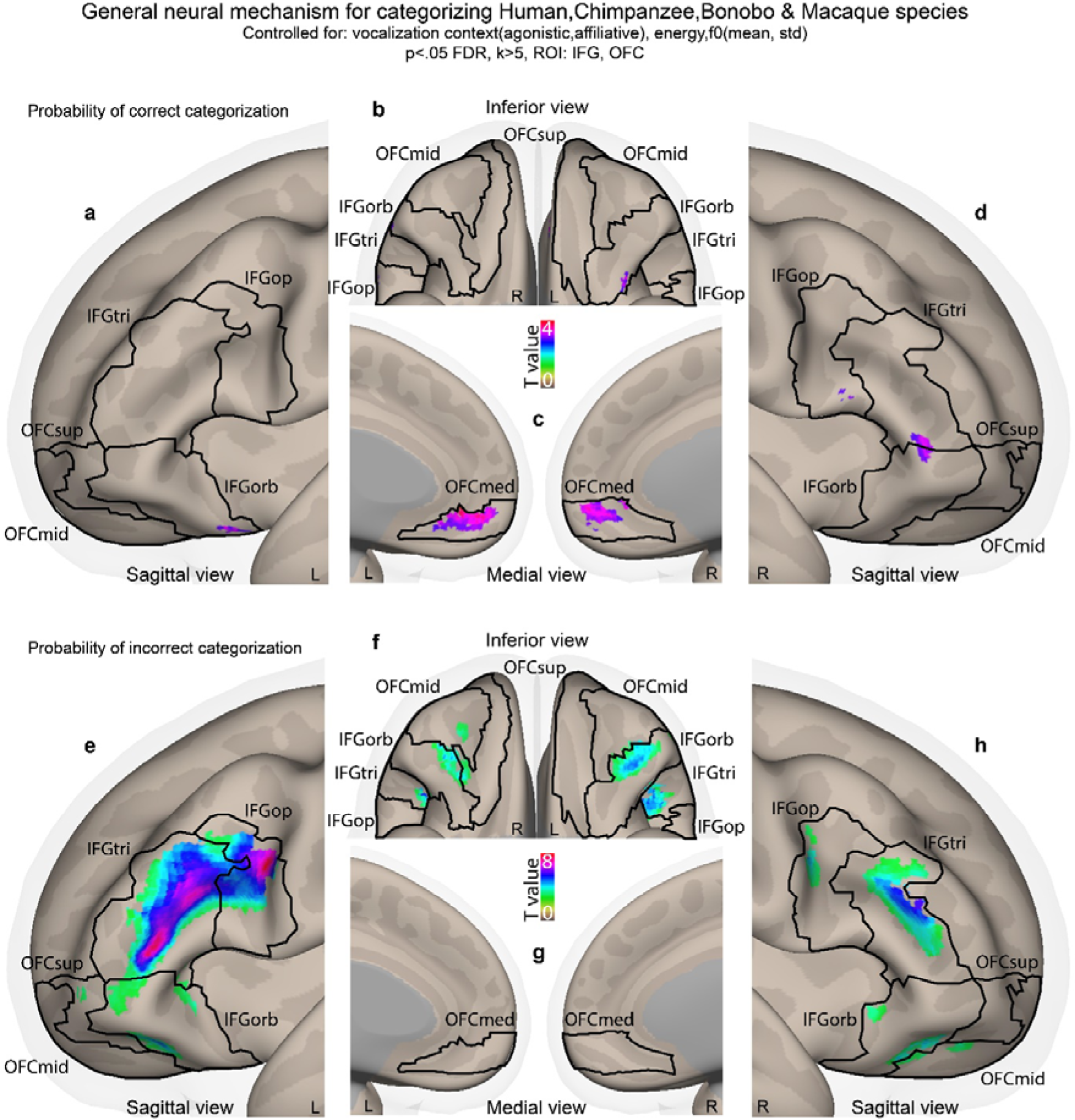
Model-based neural correlates and anti-correlates of the probability of correctly categorizing each species’ vocalizations in the IFG and OFC. **abcd,** Trial-by-trial correlates of the probability of correct species categorization. **efgh,** Trial-by-trial anti-correlates of the probability of correct species categorization. Data of this analysis were computed by using a modelling of species categorization accuracy across all participants, species (Species factor) and trials including the Affective context factor and trial-level covariates of no interest (mean and standard deviation of the vocalization energy and fundamental frequency). Subsequently, the fitted probability values of this model were used as a parametric modulator of interest for each trial’s onset, further taken as main regressor in second-level analyses. Colorbars illustrate t-value statistics. All activations thresholded at a voxelwise *p*<.05 FDR, k>5 voxels, masked by IFG and OFC regions of interest (k=9635 voxels in total). IFG: inferior frontal gyrus; tri: pars triangularis; op: pars opercularis; orb: pars orbitalis; OFC: orbitofrontal cortex; med: medial; mid: middle; sup: superior.

We then computed two conjunction analyses with each non-human primate species compared to human voice categorization. In these analyses, enhanced activations for each non-human primate species was tested while driving the conjunction but only the [(chimpanzee > human) > (bonobo > human) > (macaque > human) vocalizations] contrast showed significant results (see model 2 in the methods section). This conjunction revealed left-lateralized IFG_tri_ and IFG_op_ as well as right IFG_tri_ activity for the processing of the vocalizations (i.e., general processing displayed as white-to-pink activations, Fig.3**abd**) while enhanced activity for correctly categorized species (see model 3 in the methods section) was only found in left IFG_tri_ (i.e., correct categorization displayed as black-to-white activations, Fig.3**a**). No enhanced signal was observed in the OFC. Other combinations of non-human primate vs. human species conjunction contrasts did not yield to any above-threshold voxels. Mean and standard deviation of each vocalization’s energy and fundamental frequency were used as trial-level covariates of no interest in these statistical analyses as well. See Supplementary Table 4 for cluster coordinates.

**Figure 3.**
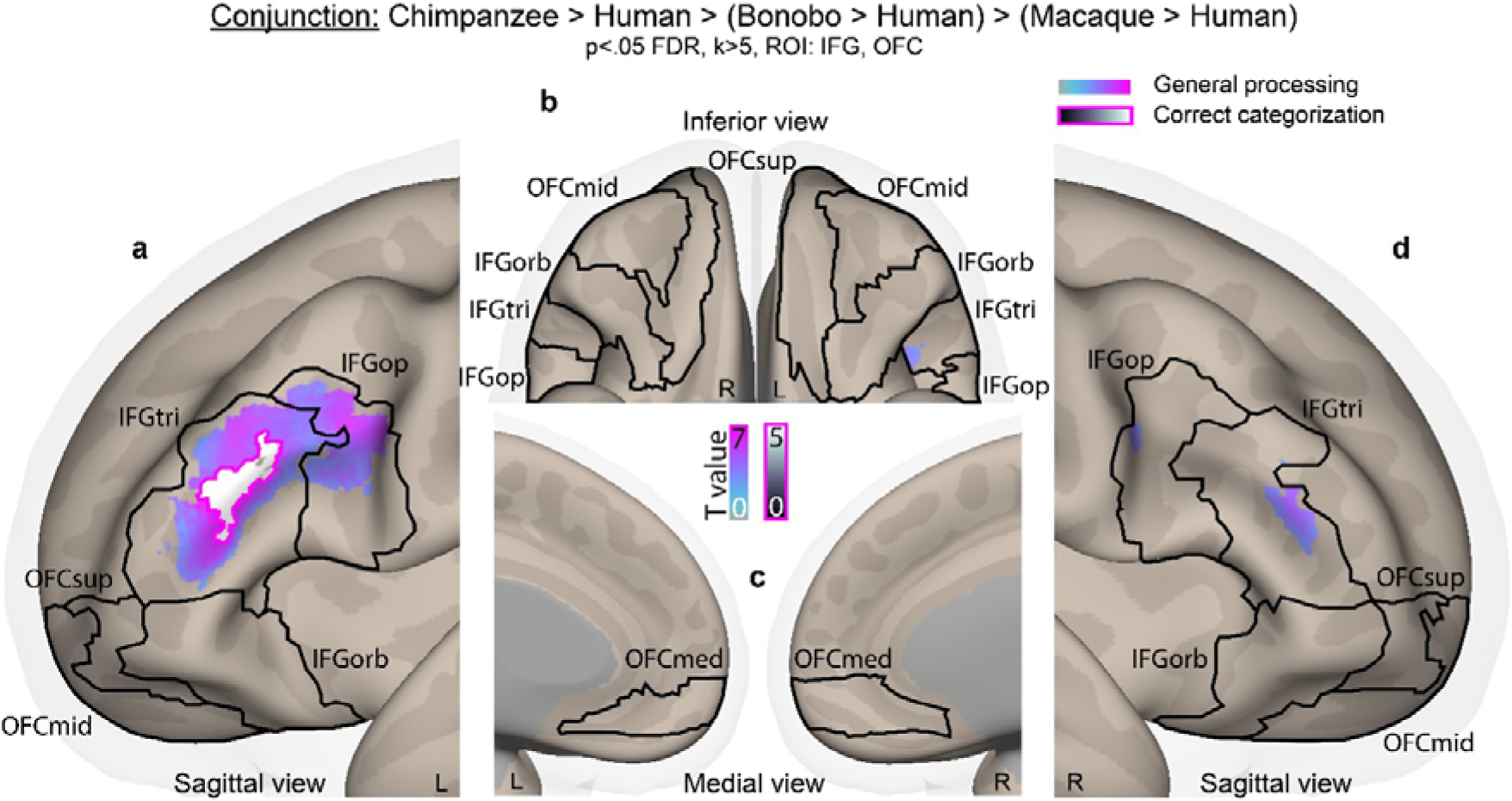
Conjunction results of the processing and correct categorization of non-human primates compared to human vocalizations in the IFG and OFC. **abcd**, Conjunction analysis for the processing (white-to-pink activations, all trials of each species) and correct species categorization (black-to-white activations with pink outline, correct trials only) of the contrasts [(chimpanzee > human) > (bonobo > human) > (macaque > human) vocalizations]. No above-threshold voxels were found for [bonobo > human] or [macaque > human] contrasts driving the conjunction analyses. For both models (species processing; correct species categorization), trial-level covariates of no interest included the mean and standard deviation of the vocalization energy and fundamental frequency. Colorbars illustrate t-value statistics. All activations thresholded at a voxelwise *p*<.05 FDR, k>5 voxels, masked by IFG and OFC regions of interest (k=9635 voxels in total). IFG: inferior frontal gyrus; tri: pars triangularis; op: pars opercularis; orb: pars orbitalis; OFC: orbitofrontal cortex; med: medial; mid: middle; sup: superior.

The comparisons between the processing (all trials) or the correct categorization (correct trials only) of the vocalizations of each non-human species compared to those of humans, and of human voices compared to the vocalizations of each non-human primate species are reported in Fig.S1 and Fig.S2, respectively. Activations specific to the processing and correct categorization of each non-human primate species or between each non-human primate species are reported in Fig.S3 and Fig.S4, respectively.

## Discussion

The present study was designed to clarify the importance of primate vocalizations processing and classification by humans as a proxy for the more general treatment of human non-verbal communication. In fact, the processing of primate vocalizations by humans might boost our understanding of non-verbal communication in today’s *Homo sapiens* and inform our reconstruction of our last common ancestor with other Hominids. To operationalize such a research objective, we investigated the behavioural and frontal brain mechanisms in modern humans underlying the vocal recognition of non-human primates’ vocalizations, considering their phylogenetic proximity to the human species. Our data highlighted the evolutionary shared history of our species with the communication system of other primates, particularly other great apes, with both the recognition and classification processes of hominid vocalizations involving the medial orbitofrontal and bilateral inferior frontal cortex, especially the *pars triangularis*.

Behavioural data emphasized the astonishing ability of human participants to categorize chimpanzee and macaque calls above chance level while it was not the case for bonobo vocalizations. Despite the close phylogenetic proximity of this species with *Homo sapiens*, the peculiar acoustical structure of bonobo calls may have prevented modern humans from recognizing them, yielding to miscategorization. Indeed, confusion matrices showed that, despite their close phylogenetic relationship to chimpanzees, bonobo vocalizations were in fact often confused for those of macaques. This peculiarity in the processing of bonobo vocalizations has also been reported in previous literature showing that human adults are highly sensitive to pitch information for discriminating the arousal of primate babies’ cries (13). In fact, human participants rated bonobo infant calls as more distressful in comparison to those of humans or chimpanzees and after further acoustical analyses, the authors found that bonobo infants’ calls had higher pitched vocalizations compared to those of human and chimpanzee babies, both species sharing more similar acoustical features (13). Such acoustical divergence of bonobos among non-human primates and especially the increased fundamental frequency of their calls may well have its origin in the shorter length of the larynx of this species (15). Interestingly in our study, chimpanzee calls were not better categorized by human participants than those of a more evolutionary distant species such as rhesus macaques. This was especially surprising considering that previous studies have mostly found a better recognition rate for chimpanzee vocalizations compared to the calls of other non-human primates (46, 51). Hence, our behavioral analysis suggests that the capacity of modern humans to accurately identify non-human primate calls could rely more heavily on acoustic similarity rather than phylogenetic proximity alone (52).

To assess the frontal brain mechanisms involved in the general processing of human voice and non-human primate vocalization recognition, we turned to computational modelling and more precisely to model-based fMRI analyses. These data pointed to a frontal network including the bilateral medial OFC for the correct recognition and categorization of human voices and non-human primate vocalizations. This first result suggests that the medial OFC has a central role in the accurate categorization of primate calls, extending previous fMRI results that showed an enhancement of activity in OFC regions—especially of the medial part—for the discrimination of emotions in human, chimpanzee, macaque, and cat vocalizations (28, 29). Other frontal brain areas underlying the probability of accurate species categorization included small clusters of the bilateral IFG_orb_ as well as the right IFG_op_ and IFG_tri_. Frontal brain areas underlying miscategorization and/or what could be labelled as ‘hesitation’ were found in large portions of the bilateral IFG, especially in the IFG_tri_ which is coherent with existing literature (53). This result is also supported by our conjunction analyses, which allowed a more concise and focused multi-contrast approach to pinpoint the specific frontal brain mechanisms that may underlie the processing as well as accurate recognition and categorization of non-human primate vocalizations compared to the human voice. These analyses revealed large, similar bilateral IFG_tri_ clusters of activity when creating a hierarchy for processing chimpanzee, bonobo and macaque vocalizations each compared to the human voice—namely, contrasting (chimpanzee > human) > (bonobo > human) > (macaque > human) at the same time. Interestingly, this conjunction contrast also revealed a difference in the hierarchy between the accurate recognition of chimpanzee vocalizations and those of bonobos and macaques when each was compared to the human voice in a subpart of the left IFG_tri_. In the literature, the IFG_tri_ was repeatedly involved when human participants have to make a decision based on vocal signals, for instance when human participants discriminate or categorize human vocal emotion (37, 50) or when they implicitly or explicitly process human emotional voices (7). Causal links between IFG_tri_ functioning and emotion recognition were also revealed in the literature (54). Noteworthy is the fact that activity in the left IFG_tri_ also translates a level of uncertainty when decisional processes occur (53, 55–57). Combining our behavioural and neuroimaging results, it seems extremely plausible that decisional uncertainty was peaking when human participants were categorizing non-human primate vocalizations. We therefore think that according to our results and to the existing literature, activity in the IFG_tri_ might underlie both categorization uncertainty as revealed by anti-correlates of the probability of accurate species recognition in the model-based neuroimaging results, as well as decisional processes that lead to an accurate species recognition in a very concise region of the left IFG_tri_, as highlighted by our conjunction results. Therefore, OFC and IFG brain areas—especially IFGtri activity—may be the necessary biological and functional ‘hardware’ allowing *Homo sapiens* to classify and integrate non-verbal communication, be it human voice or other primate vocalizations. Such ability may then have been co-opted for the treatment of the proto-language used by our human ancestors, although precisely how close this proto-language was from to today’s non-human primates vocalizations remains an open question, with a question mark regarding a potential interaction with between-species acoustical differences (52). We contend that the common activity we evidenced in the OFC and IFG across stimuli suggests that such proto-language may have shared acoustic properties with non-human primate vocalizations.

Even though our results shed new light on the origins of human non-verbal communication, our study suffered from several limitations. Such limitations include the absence of higher temporal resolution, the absence of other great apes as stimuli (for instance gorillas and/or orangutans) and the absence of highly specific acoustic distance assessment. Also, the time-course of such frontal mechanisms underlying non-verbal communication in humans should be investigated.

To conclude, the present study revealed the existence of distinct mechanisms at play in the human frontal cortex for the integration and further categorization of non-human primate vocalizations. In fact, computational modelling of the fMRI data allowed a clear distinction between accurate categorization on the one hand and miscategorization processes on the other hand, as revealed by bilateral medial OFC and left IFG_tri_, respectively. Combined with our behavioural data, evidence converges toward a preserved ability of *Homo sapiens* to recognize the calls of species close to his ancestors that were most likely very similar to those of today’s non-human primate vocalizations. Natural selection might also have led to specific anatomical adaptations in some non-human primates across time, and the shorter larynx of bonobos leading to the significantly increased fundamental frequency of their calls is a well-fitting example, especially in the context of the present study. Therefore, a combination of phylogenetic and acoustical proximity may be necessary for humans to recognize and successfully categorize the calls of non-human primates, including those of great apes. Overall, the present study substantially contributes to the understanding of primate calls’ recognition by modern humans as a proxy for more general non-verbal communication mechanisms in *Homo sapiens*.

## Material and Methods

### Experimental procedure and paradigm

Participants laid comfortably in a 3T scanner and heard a total of seventy–two stimuli in pseudo-randomized order through MRI-compatible earphones (Sensimetrics Corporation, Gloucester, MA, USA) at a sound pressure level of 70dB. Participants were instructed to identify the species that expressed the vocalizations using a response box (Current Designs, Inc., Philadelphia, PA, USA). For instance, the instructions could be “Human – press 1, Chimpanzee – press 2, Bonobo – press 3 or Macaque – press 4”. There were four different mappings of the keys that were randomly assigned across participants. In a 3 – 5 second interval (jittering of 400 ms) after each stimulus, participants were asked to categorize the species. If the participant did not respond during this interval—namely a missed trial, the next stimulus followed automatically and the value of the response was ‘0’ while ‘1’ coded a correct response, ‘2’ an incorrect response (Fig.1**a**).

### Stimuli

Seventy-two vocalizations of four primate species (human, chimpanzee, bonobo and rhesus macaque) were used in this study. The eighteen human voices were expressed by two male and two female actors, obtained from a non-verbal (affective vocal burst) validated set of Belin and collaborators (58). Non-linguistic vocalizations were selected as human voice stimuli due to their acoustic and perceptual proximity with the other mammalian calls, non-human primates included (44). The eighteen selected chimpanzee, bonobo and rhesus macaque vocalizations contained single calls or two call sequences produced by 6 to 8 different individuals in their natural environment. For each non-human primate species, the stimuli were either produced in an affiliative context or produced in an agonistic context (threating or distressful). All vocal stimuli were standardized to 750 milliseconds using PRAAT (www.praat.org) but were not normalized in order to preserve the ‘naturality’ of the sounds (59).

### Participants

Twenty-five right-handed, healthy, either native or highly proficient French-speaking participants took part in the study. One participant was excluded because he had no correct response at all and may have fallen asleep, while another participant was excluded due to incomplete scanning and technical issues, leaving us with twenty-three participants (10 female, 13 male, mean age 24.65 years, SD 3.66). All participants were naive to the experimental design and study, had normal or corrected-to-normal vision, normal hearing and no history of psychiatric or neurologic incidents. Participants gave written informed consent for their participation in accordance with ethical and data security guidelines of the University of Geneva. The study was approved by the Ethics Cantonal Commission for Research of the Canton of Geneva, Switzerland (CCER) and was conducted according to the Declaration of Helsinki.

### Statistical analysis

Behavioural data were analysed using R studio software (60). In order to test our hypotheses regarding the impact of phylogenetic distance on species categorization, we performed mixed-effects models analyses of the ‘lme4’ package (61) on accuracy and confusion (one accuracy column: ‘1’, correct response; ‘0’, incorrect response; four confusion columns: ‘1’, confusion with the species, ‘0’ no confusion with the species, for each of the four species column) and reaction times data (for correct, incorrect trials and specific confusion for each species) with Species (human, chimpanzee, bonobo, macaque) and Affective context (affiliative and agonistic) as interacting fixed factors, four covariates (mean energy of each species, mean fundamental frequency of each species, energy standard deviation of each species, fundamental frequency standard deviation of each species) and the participants’ and the stimuli identity as random factors (random intercept and random slope, respectively). The inclusion of the sex and age of the participants for which we had no specific hypothesis in data modelling did not yield to a significantly better model neither for accuracy (χ^2^(2)=.81, *p*=.66) nor for reaction times data (χ^2^(2)=.01, *p*=1) and these random effects were therefore discarded from the analyses. The same model was used to analyse accuracy, confusion and reaction times data, except that a logistic regression (generalized linear mixed-effects models, ‘glmer’) was used for accuracy and a linear regression (linear mixed-effects models, ‘lmer’) for specific pairs of species confusion (four models in total) and for reaction times. Then, we used specific contrasts computed using the ‘phia’ package (62) to compare between-species categorization, testing our hypothesis of an impact of phylogenetic proximity on species categorization. Contrasts therefore included human vs. non-human primate planned comparisons: [human > chimpanzee, bonobo, macaque vocalizations], [human > chimpanzee vocalizations], [human > bonobo vocalizations], [human > macaque vocalizations]; non-human primate vs. human comparisons: [chimpanzee > human vocalizations], [bonobo > human vocalizations], [macaque > human vocalizations]. These inverse contrasts actually yield to a sign-reversal of the difference estimate of the original contrast for accuracy data. For confusion data, contrasts were computed to uncover any difference between one human or non-human primate species compared to the remaining species: [Confusion for human vocalizations: chimpanzee response ≠ bonobo response ≠ macaque response], [Confusion for chimpanzee vocalizations: human response ≠ bonobo response ≠ macaque response], [Confusion for bonobo vocalizations: human response ≠ chimpanzee response ≠ macaque response], [Confusion for macaque vocalizations: human response ≠ chimpanzee response ≠ bonobo response]. All contrasts were corrected for multiple comparisons using a Bonferroni adjustment method implemented in the ‘phia’ package (adjustment=”Bonf”) and *p*-value is accordingly reported as ‘*p_Bonf_*’. Reaction times data were analysed using the same contrasts and are reported in Supplementary Material.

### Image acquisition

Structural and functional brain imaging data were acquired by using a 3T scanner Siemens Trio, Erlangen, Germany with a 32-channel coil. A 3D GR\IR magnetization-prepared rapid acquisition gradient echo sequence was used to acquire high-resolution (0.35 × 0.35 × 0.7 mm^3^) T1-weighted structural images (TR = 2400 ms, TE = 2.29 ms). Functional images were acquired by using fast fMRI, with a multislice echo planar imaging sequence 79 transversal slices in descending order, slice thickness 3 mm, TR = 650 ms, TE = 30 ms, field of view = 205 × 205 mm2, 64 × 64 matrix, flip angle = 50 degrees, bandwidth 1562 Hz/Px.

### Wholebrain image analysis

Functional images were analysed with Statistical Parametric Mapping software (SPM12, Wellcome Trust Centre for Neuroimaging, London, UK). Preprocessing steps included realignment to the first volume of the time series, slice timing, normalization into the Montreal Neurological Institute (MNI) space (63) using the DARTEL toolbox (64) and spatial smoothing with an isotropic Gaussian filter of 8 mm full width at half maximum. To remove low-frequency components, we used a high-pass filter with a cut-off frequency of 128s. Three general linear models were used to compute first-level statistics.

#### Model 1: model-based categorization probability of each species, including all trials

In this model, the probability of correctly categorizing each species in each production context was modelled using the ‘fitted’ function of the ‘stats’ package (65) to obtain the fitted probability values from the behavioural accuracy data described in the behavioural data analysis section. This modelling was performed by using each trial (N=72) of each fMRI session (N=1) of each participant (N=23) with a binary ‘1’ and ‘0’ coding for the correct and incorrect species categorization, respectively, as dependent variable. As mentioned above, the model included fixed effects, namely the interaction between the Species factor (human, chimpanzee, bonobo, macaque) and the Affective context factor (affiliative, agonistic) and four covariates of interest (in this order: mean energy of each species, mean fundamental frequency of each species, energy standard deviation of each species, fundamental frequency standard deviation of each species). Random effects included the participants’ and the stimuli identity (random intercept and random slope, respectively). The sample-specific matrix (participants per trials=23*72=1656 lines) was then cut to the corresponding participant-specific matrix (72 lines). Model 1 therefore included two 72-lines regressors, namely the onsets of each trial in its order of appearance—modelled by using a boxcar function and convolved with the hemodynamic response function time-locked to the onset of each stimulus—and the corresponding modelled probability of correctly categorizing each species as a covariate of interest. Six motion parameters were included as regressors of no interest to account for movement in the data and our design matrix therefore included a total of 8 columns plus the constant term. The regressor of interest (modelled probability of correctly categorizing each species) was used to compute a simple contrast for each participant, leading to separate main effect of the probability of correctly categorizing each species. This simple contrast was then taken to a one-sample t-test second-level analysis in which variance was set to ‘unequal’.

#### Model 2: processing of all species, including all trials

In this model, each event was modelled by using a boxcar function and was convolved with the hemodynamic response function, time-locked to the onset of each stimulus. Separate regressors were created for all trials of each species (Species factor: human, chimpanzee, bonobo, macaque) and production context (Affective context factor: affiliative, agonistic) and four covariates each (mean energy of each species, mean fundamental frequency of each species, energy standard deviation of each species, fundamental frequency standard deviation of each species) for a total of 47 regressors. The means and standard deviations of the fundamental frequency and the energy of the vocalizations were extracted using the extended Geneva Acoustic parameters set, which is defined as the optimal acoustic indicators related to human voice analysis (GeMAPS (66)). This set of acoustical parameters was selected based on i) their potential to index affective physiological changes in voice production, ii) their proven value in former studies as well as their automatic extractability, and iii) their theoretical significance.

Six motion parameters were included as regressors of no interest to account for movement in the data and our design matrix therefore included a total of 66 columns plus the constant term. The species regressors were used to compute simple contrasts for each participant, leading to separate main effects of human, chimpanzee, bonobo and macaque vocalizations for each production context (12 simple contrasts). Covariates were set to zero in order to model them as no-interest regressors. These simple contrasts were then taken to a flexible factorial second-level analysis in which there were three factors: the Participants factor (data independence set to ‘yes’, variance set to ‘unequal’), the Species factor and the Affective context factor (for both factors: data independence set to ‘no’, variance set to ‘unequal’). For this first model, the main effect of the Species factor was computed in the flexible factorial modelling while the Affective context factor was modelled but excluded from the results because we did not have any hypothesis regarding this factor and because each context modality contained only 6 trials per species.

#### Model 3: correct categorization of each species, including only correct trials

Separate regressors were created for each correctly categorized species (Correct Species factor: human hits, chimpanzee hits, bonobo hits, macaque hits vocalizations) for each production context (Affective context factor: affiliative, agonistic) and four covariates each (mean energy of each species, mean fundamental frequency of each species, energy standard deviation of each species, fundamental frequency standard deviation of each species) as well as a regressor containing a concatenation of categorization errors across species for a total of 44 regressors. Regressors were modelled by using a boxcar function and convolved with the hemodynamic response function, time-locked to the onset of each stimulus. Six motion parameters were included as regressors of no interest to account for movement in the data and our design matrix therefore included a total of 51 columns plus the constant term. The regressors of interest (Correct Species factor) were used to compute 8 simple contrasts for each participant, leading to separate main effects of human hits, chimpanzee hits, bonobo hits and macaque hits vocalizations for each production context. Covariates were set to zero in order to model them as no-interest regressors. These simple contrasts were then taken to a flexible factorial second-level analysis in which there were three factors: the Participants factor (independence set to ‘yes’, variance set to ‘unequal’), the Correct Species factor and the Affective context factor (for both factors: independence set to ‘no’, variance set to ‘unequal’). For this third model, the main effect of the Correct Species factor was computed in the flexible factorial modelling while the Affective context factor was modelled but excluded from the results because we did not have any hypothesis regarding this factor and because each modality contained only 6 trials per species.

For all three models and to be consistent in our analyses, neuroimaging activations were masked by a binary image including all subparts of the IFG (*pars opercularis*, *triangularis* and *orbitalis*) and OFC (superior, middle and medial) for a total of twelve subregions, 9635 voxels and thresholded in SPM12 by using a voxel-wise false discovery rate (FDR) correction at *p*<.05 and an arbitrary cluster extent of k>5 voxels to remove extremely small clusters of activity. Region masks were extracted using the ‘automated anatomical labelling’ (‘aal’) atlas (67) implemented in the ‘WFU_Pickatlas’ toolbox (https://www.nitrc.org/projects/wfu_pickatlas/). Contrast activations were rendered on semi-inflated brains from the CONN toolbox (68) with black outlines delineating each of our IFG-OFC regions of interest.

## Acknowledgements

We warmly thank Prof. Katie Slocombe and Dr. Zanna Clay for providing the non-human primates vocalizations as well as Prof. Daphne Bavelier for her valuable advice on the experimental task. We would like also to acknowledge the staff of the Brain and Behaviour Laboratory at The University of Geneva where all data were acquired.

## References

1. Tracy JL, Randles D, Steckler CM. The nonverbal communication of emotions. Current opinion in behavioral sciences. 2015;3:25–30.

2. Leger DW. Contextual sources of information and responses to animal communication signals. Psychol Bull. 1993;113(2):295.

3. Marler P. Animal Communication Signals: We are beginning to understand how the structure of animal signals relates to the function they serve. Science. 1967;157(3790):769-74.

4. Gagneux P, Varki A. Genetic differences between humans and great apes. Molecular phylogenetics and evolution. 2001;18(1):2–13.

5. Prado-Martinez J, Sudmant PH, Kidd JM, Li H, Kelley JL, Lorente-Galdos B, et al. Great ape genetic diversity and population history. Nature. 2013;499(7459):471-5.

6. Binder JR, Liebenthal E, Possing ET, Medler DA, Ward BD. Neural correlates of sensory and decision processes in auditory object identification. Nature Neuroscience. 2004;7(3):295–301.

7. Frühholz S, Ceravolo L, Grandjean D. Specific brain networks during explicit and implicit decoding of emotional prosody. Cereb Cortex. 2012;22(5):1107–17.

8. Grandjean D. Brain Networks of Emotional Prosody Processing. Emotion Review. 2020.

9. Murray EA, Izquierdo A. Orbitofrontal Cortex and Amygdala Contributions to Affect and Action in Primates. Annals of the New York Academy of Sciences. 2007;1121(1):273–96.

10. Damasio AR, Anderson SW. The frontal lobes. 2003.

11. Semendeferi K, Lu A, Schenker N, Damasio H. Humans and great apes share a large frontal cortex. Nature Neuroscience. 2002;5(3):272–6.

12. Perelman P, Johnson WE, Roos C, Seuánez HN, Horvath JE, Moreira MAM, et al. A Molecular Phylogeny of Living Primates. PLOS Genetics. 2011;7(3):e1001342.

13. Kelly T, Reby D, Levréro F, Keenan S, Gustafsson E, Koutseff A, et al. Adult human perception of distress in the cries of bonobo, chimpanzee, and human infants. Biological Journal of the Linnean Society. 2017;120(4):919–30.

14. Graham KE, Hobaiter C. Towards a great ape dictionary: Inexperienced humans understand common nonhuman ape gestures. PLOS Biology. 2023;21(1):e3001939.

15. Grawunder S, Crockford C, Clay Z, Kalan AK, Stevens JMG, Stoessel A, et al. Higher fundamental frequency in bonobos is explained by larynx morphology. Current biology: CB. 2018;28(20):R1188–R9.

16. Gruber T, Clay Z. A Comparison Between Bonobos and Chimpanzees: A Review and Update. Evolutionary Anthropology: Issues, News, and Reviews. 2016;25(5):239–52.

17. Hare B, Wobber V, Wrangham R. The self-domestication hypothesis: evolution of bonobo psychology is due to selection against aggression. Animal Behaviour. 2012;83(3):573–85.

18. Staes N, Smaers JB, Kunkle AE, Hopkins WD, Bradley BJ, Sherwood CC. Evolutionary divergence of neuroanatomical organization and related genes in chimpanzees and bonobos. Cortex. 2018.

19. Herculano-Houzel S. The human brain in numbers: a linearly scaled-up primate brain. Frontiers in Human Neuroscience. 2009;3.

20. Barbas H. Connections underlying the synthesis of cognition, memory, and emotion in primate prefrontal cortices. Brain Res Bull. 2000;52(5):319–30.

21. Barbas H, Zikopoulos B, Timbie C. Sensory Pathways and Emotional Context for Action in Primate Prefrontal Cortex. Biological Psychiatry. 2011;69(12):1133–9.

22. Davidson RJ. Anterior cerebral asymmetry and the nature of emotion. Brain and Cognition. 1992;20(1):125–51.

23. Frühholz S, Grandjean D. Processing of emotional vocalizations in bilateral inferior frontal cortex. Neuroscience and Biobehavioral Reviews. 2013;37(10 Pt 2):2847-55.

24. Kambara T, Brown EC, Silverstein BH, Nakai Y, Asano E. Neural dynamics of verbal working memory in auditory description naming. Scientific Reports. 2018;8(1):15868.

25. LeDoux J. Rethinking the emotional brain. Neuron. 2012;73(4):653–76.

26. Rolls ET. Convergence of sensory systems in the orbitofrontal cortex in primates and brain design for emotion. The Anatomical Record Part A: Discoveries in Molecular, Cellular, and Evolutionary Biology. 2004;281A(1):1212-25.

27. Sander D, Grandjean D, Scherer KR. A systems approach to appraisal mechanisms in emotion. Neural Networks. 2005;18(4):317–52.

28. Belin P, Fecteau S, Charest I, Nicastro N, Hauser MD, Armony JL. Human cerebral response to animal affective vocalizations. Proc Biol Sci. 2008;275(1634):473-81.

29. Fritz T, Mueller K, Guha A, Gouws A, Levita L, Andrews TJ, et al. Human behavioural discrimination of human, chimpanzee and macaque affective vocalisations is reflected by the neural response in the superior temporal sulcus. Neuropsychologia. 2018;111:145–50.

30. Hsu C-C, Rolls ET, Huang C-C, Chong ST, Zac Lo C-Y, Feng J, et al. Connections of the Human Orbitofrontal Cortex and Inferior Frontal Gyrus. Cerebral Cortex. 2020;30(11):5830–43.

31. Petrides M, Pandya DN. Comparative cytoarchitectonic analysis of the human and the macaque ventrolateral prefrontal cortex and corticocortical connection patterns in the monkey. Eur J Neurosci. 2002;16(2):291–310.

32. Guenther FH, Tourville JA, Bohland JW. Speech Production. In: Toga AW, editor. Brain Mapping. Waltham: Academic Press; 2015. p. 435-44.

33. Brück C, Kreifelts B, Kaza E, Lotze M, Wildgruber D. Impact of personality on the cerebral processing of emotional prosody. NeuroImage. 2011;58(1):259–68.

34. Hampshire A, Chamberlain SR, Monti MM, Duncan J, Owen AM. The role of the right inferior frontal gyrus: inhibition and attentional control. NeuroImage. 2010;50(3-3):1313–9.

35. Tops M, Boksem MAS. A Potential Role of the Inferior Frontal Gyrus and Anterior Insula in Cognitive Control, Brain Rhythms, and Event-Related Potentials. Frontiers in Psychology. 2011;2.

36. Ethofer T, Anders S, Erb M, Herbert C, Wiethoff S, Kissler J, et al. Cerebral pathways in processing of affective prosody: a dynamic causal modeling study. NeuroImage. 2006;30(2):580–7.

37. Gruber T, Debracque C, Ceravolo L, Igloi K, Marin Bosch B, Frühholz S, et al. Human Discrimination and Categorization of Emotions in Voices: A Functional Near-Infrared Spectroscopy (fNIRS) Study. Frontiers in Neuroscience. 2020;14.

38. Zhang D, Zhou Y, Yuan J. Speech Prosodies of Different Emotional Categories Activate Different Brain Regions in Adult Cortex: an fNIRS Study. Scientific Reports. 2018;8(1):218.

39. Belyk M, Brown S, Lim J, Kotz SA. Convergence of semantics and emotional expression within the IFG pars orbitalis. NeuroImage. 2017;156:240–8.

40. Schirmer A, Kotz SA. Beyond the right hemisphere: brain mechanisms mediating vocal emotional processing. Trends in Cognitive Sciences. 2006;10(1):24–30.

41. Stout D, Chaminade T. Stone tools, language and the brain in human evolution. Philosophical Transactions of the Royal Society B: Biological Sciences. 2012;367(1585):75-87.

42. Ethofer T, Bretscher J, Gschwind M, Kreifelts B, Wildgruber D, Vuilleumier P. Emotional voice areas: anatomic location, functional properties, and structural connections revealed by combined fMRI/DTI. Cereb Cortex. 2012;22(1):191–200.

43. Gruber T, Grandjean DM. A comparative neurological approach to emotional expressions in primate vocalizations. Neuroscience and Biobehavioral Reviews. 2017;73:182–90.

44. Anikin A, Bååth R, Persson T. Human non-linguistic vocal repertoire: Call types and their meaning. Journal of Nonverbal Behavior. 2018;42(1):53–80.

45. Belin P. Voice processing in human and non-human primates. Philosophical Transactions of the Royal Society B: Biological Sciences. 2006;361(1476):2091-107.

46. Filippi P, Congdon JV, Hoang J, Bowling DL, Reber SA, Pašukonis A, et al. Humans recognize emotional arousal in vocalizations across all classes of terrestrial vertebrates: evidence for acoustic universals. Proceedings of the Royal Society B: Biological Sciences. 2017;284(1859):20170990.

47. Kamiloğlu RG, Slocombe KE, Haun DBM, Sauter DA. Human listeners’ perception of behavioural context and core affect dimensions in chimpanzee vocalizations. Proceedings of the Royal Society B: Biological Sciences. 2020;287(1929):20201148.

48. Scheumann M, Hasting AS, Kotz SA, Zimmermann E. The voice of emotion across species: how do human listeners recognize animals’ affective states? PLOS ONE. 2014;9(3):e91192.

49. Scheumann M, Hasting AS, Zimmermann E, Kotz SA. Human Novelty Response to Emotional Animal Vocalizations: Effects of Phylogeny and Familiarity. Frontiers in Behavioral Neuroscience. 2017;11.

50. Dricu M, Ceravolo L, Grandjean D, Frühholz S. Biased and unbiased perceptual decision-making on vocal emotions. Scientific Reports. 2017;7(1):16274.

51. Linnankoski I, Laakso M, Aulanko R, Leinonen L. Recognition of emotions in macaque vocalizations by children and adults. Language & Communication. 1994;14(2):183–92.

52. Debracque C, Clay Z, Grandjean D, Gruber T. Humans recognize affective cues in primate vocalizations: Acoustic and phylogenetic perspectives. 2022:Preprint at https://www.biorxiv.org/content/10.1101/2022.01.26.477864v1.

53. Moss H, Abdallah S, Fletcher P, Bright P, Pilgrim L, Acres K, et al. Selecting among competing alternatives: selection and retrieval in the left inferior frontal gyrus. Cerebral Cortex. 2005;15(11):1723–35.

54. Keuken MC, Hardie A, Dorn BT, Dev S, Paulus MP, Jonas KJ, et al. The role of the left inferior frontal gyrus in social perception: an rTMS study. Brain Res. 2011;1383:196–205.

55. Ceravolo L, Moisa M, Grandjean D, Ruff C, Fruhholz S. Disrupting inferior frontal cortex activity alters affect decoding efficiency from clear but not from ambiguous affective speech. 2021:Preprint at https://www.biorxiv.org/content/10.1101/2021.12.15.472758v2.

56. Nastase S, Iacovella V, Hasson U. Uncertainty in visual and auditory series is coded by modality[general and modality[specific neural systems. Human Brain Mapping. 2014;35(4):1111–28.

57. Toelch U, Bach DR, Dolan RJ. The neural underpinnings of an optimal exploitation of social information under uncertainty. Social Cognitive and Affective Neuroscience. 2014;9(11):1746–53.

58. Belin P, Fillion-Bilodeau S, Gosselin F. The Montreal Affective Voices: A validated set of nonverbal affect bursts for research on auditory affective processing. ResearchGate. 2008.

59. Ferdenzi C, Patel S, Mehu-Blantar I, Khidasheli M, Sander D, Delplanque S. Voice attractiveness: Influence of stimulus duration and type. Behavior Research Methods. 2013;45(2):405–13.

60. Team R. RStudio: Integrated Development for R. RStudio. Boston MA: RStudio, Inc.; 2020.

61. Bates D, Mächler M, Bolker B, Walker S. Fitting Linear Mixed-Effects Models Using lme4. Journal of Statistical Software. 2015;67(1):1–48.

62. De Rosario-Martinez H, Fox J, Team R, De Rosario-Martinez MH. Package ‘Phia’. 2015;Retrieved 1.

63. Collins DL, Neelin P, Peters TM, Evans AC. Automatic 3D intersubject registration of MR volumetric data in standardized Talairach space. J Comput Assist Tomogr. 1994;18(2):192–205.

64. Ashburner J. A fast diffeomorphic image registration algorithm. NeuroImage. 2007;38(1):95–113.

65. Team RC. R: A language and environment for statistical computing. 2013.

66. Eyben F, Scherer K, Schuller B, Sundberg J, André E, Busso C, et al. The Geneva Minimalistic Acoustic Parameter Set (GeMAPS) for Voice Research and Affective Computing. IEEE transactions on affective computing. 2016;7(2):190–202.

67. Tzourio-Mazoyer N, Landeau B, Papathanassiou D, Crivello F, Etard O, Delcroix N, et al. Automated Anatomical Labeling of Activations in SPM Using a Macroscopic Anatomical Parcellation of the MNI MRI Single-Subject Brain. NeuroImage. 2002;15(1):273–89.

68. Whitfield-Gabrieli S, Nieto-Castanon A. Conn: a functional connectivity toolbox for correlated and anticorrelated brain networks. Brain Connect. 2012;2(3):125–41.

