## Supplementary figures and tables for "Frontal mechanisms underlying primate calls recognition by humans"

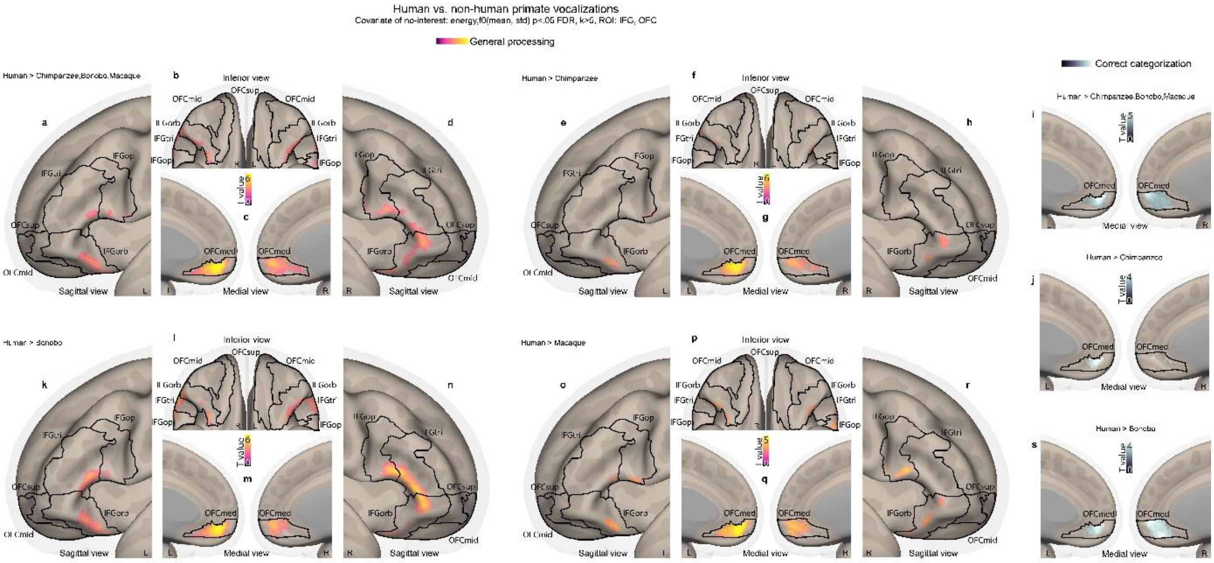

**Fig.S1: Neural activations of the processing and correct categorization of human compared to non-human primates vocalizations in the IFG and OFC.** Enhanced activations for the processing (all trials of each species) of [human > chimpanzee, bonobo, macaque vocalizations, **abcd**], [human > chimpanzee vocalizations, **efgh**], [human > bonobo vocalizations, **klmn**] and [human > macaque vocalizations, **opqr**]. Enhanced activations for the correct categorization (correct trials only) of [human > chimpanzee, bonobo, macaque vocalizations, **i**], [human > chimpanzee vocalizations, **j**] and [human > bonobo vocalizations, **s**]. No above-threshold voxels were found for correctly categorized [human > macaque vocalizations]. For both models (species processing; correct species categorization), trial-level covariates of no interest included the mean and standard deviation of the vocalization energy and fundamental frequency. Colorbars illustrate t-value statistics, with purple-to-yellow bars and activations used for contrasts of species processing while black-to-white was used for correct species categorization. All activations thresholded at a voxelwise p<.05 FDR, k>5 voxels, masked by IFG and OFC regions of interest (k=9635 voxels in total). IFG: inferior frontal gyrus; tri: pars triangularis; op: pars opercularis; orb: pars orbitalis; OFC: orbitofrontal cortex; med: medial; mid: middle; sup: superior.

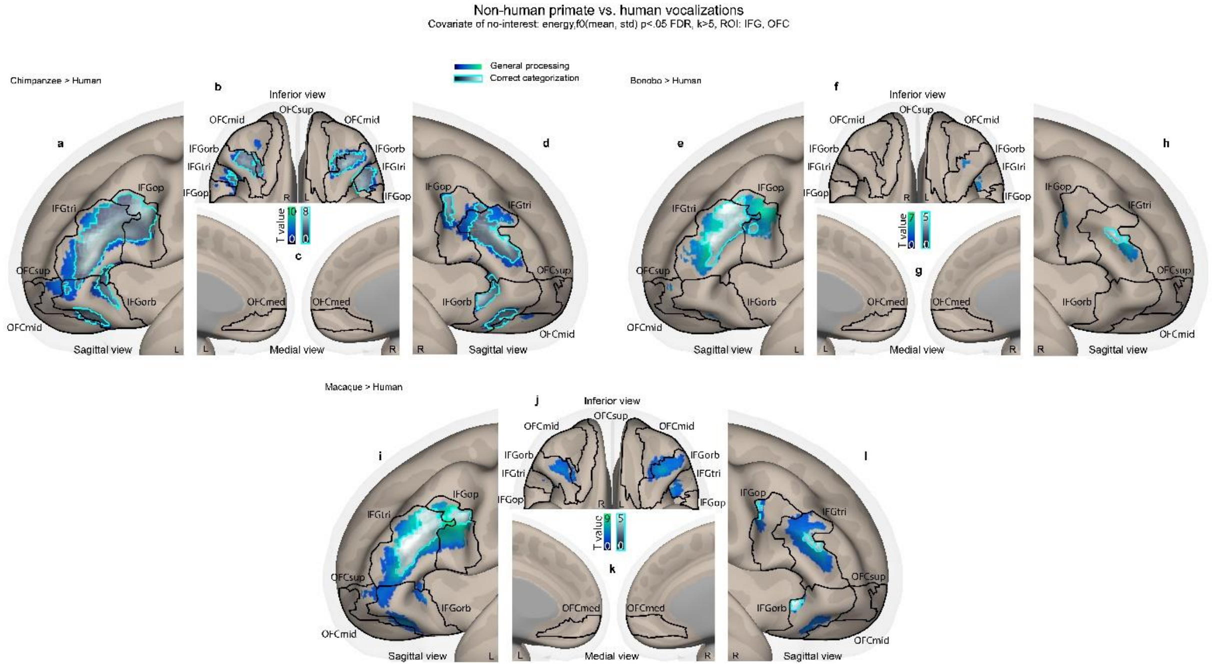

**Fig.S2: Neural activations of the processing and correct categorization of non-human primates compared to human vocalizations in the IFG and OFC.** Enhanced activations for the processing (blue-to-green activations, all trials of each species) and correct species categorization (black-to-white activations with teal outline, correct trials only) of [chimpanzee > human vocalizations, **abcd**], [bonobo > human vocalizations, **efgh**] and [macaque > human vocalizations, **ijkl**]. For both analyses (species processing; correct species categorization), trial-level covariates of no interest included the mean and standard deviation of the vocalization energy and fundamental frequency. Colorbars illustrate t-value statistics. All activations thresholded at a voxelwise $p<.05$ FDR, k>5 voxels, masked by IFG and OFC regions of interest (k=9635 voxels in total). IFG: inferior frontal gyrus; tri: pars triangularis; op: pars opercularis; orb: pars orbitalis; OFC: orbitofrontal cortex; med: medial; mid: middle; sup: superior.

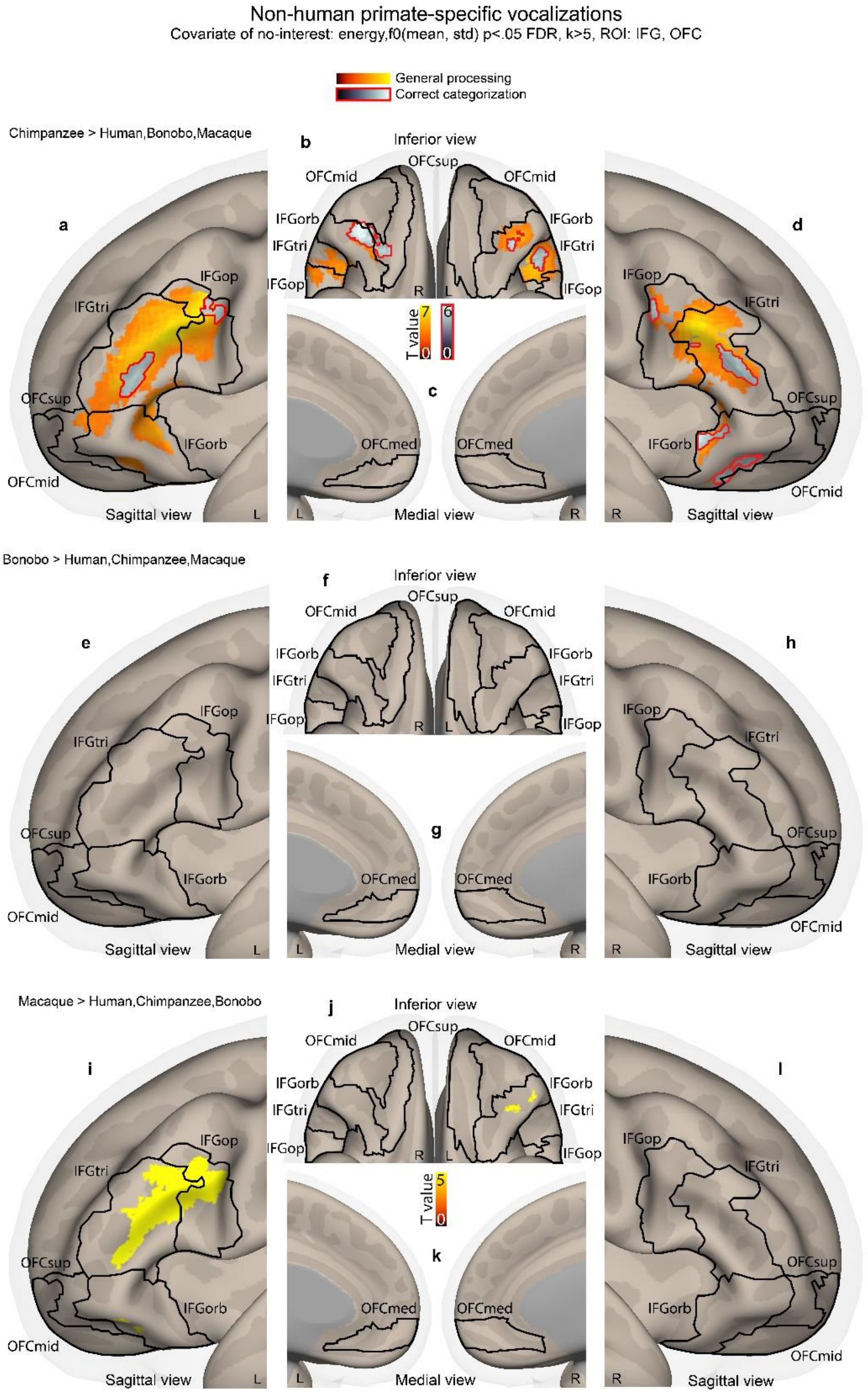

**Fig.S3: Brain activations specific to each non-human primate species within the IFG and OFC. abcd**: Processing and correct responses for Chimpanzee > Bonobo, Macaque, Human vocalizations. **efgh**: Bonobo > Chimpanzee, Macaque, Human vocalizations processing. **ijkl**: Macaque > Chimpanzee, Bonobo, Human vocalizations processing. For both models (species processing; correct species categorization), trial-level covariates of no interest included the mean and standard deviation of the vocalization energy and fundamental frequency. Colorbars illustrate t-value statistics, with red-to-yellow bars and activations used for contrasts of species processing while black-to-white with red outline was used for correct species categorization. All activations thresholded at a voxelwise $p<.05$ FDR, $k>5$ voxels, masked by IFG and OFC regions of interest ($k=9635$ voxels in total). IFG: inferior frontal gyrus; tri: *pars triangularis*; op: *pars opercularis*; orb: *pars orbitalis*; OFC: orbitofrontal cortex; med: medial; mid: middle; sup: superior.

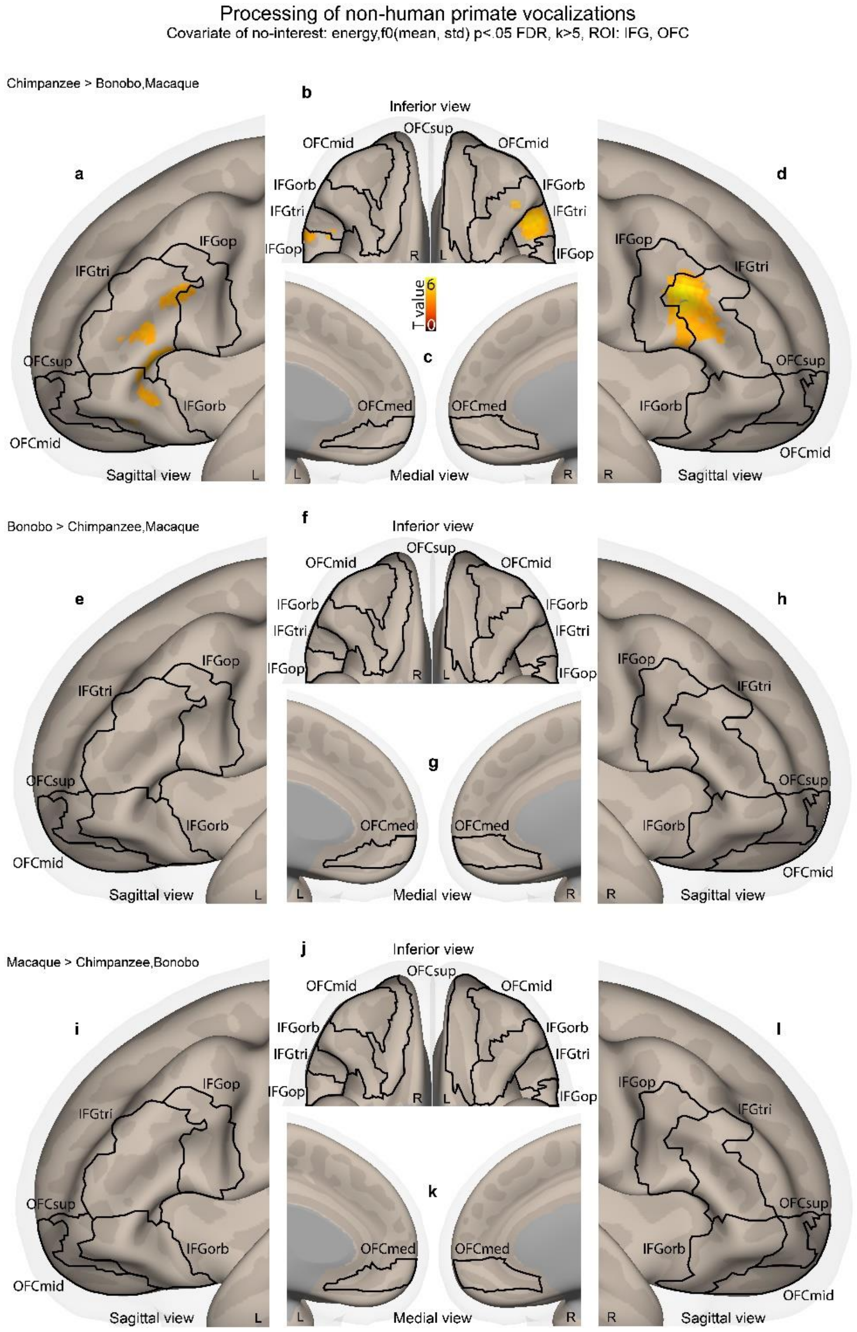

**Fig.S4: Brain activations of non-human primate vocalizations processing within the IFG and OFC. abcd**: Processing for Chimpanzee > Bonobo, Macaque vocalizations. **efgh**: Bonobo > Chimpanzee, Macaque vocalizations processing. **ijkl**: Macaque > Chimpanzee, Bonobo vocalizations processing. Trial-level covariates of no interest included the mean and standard deviation of the vocalization energy and fundamental frequency. Colorbars illustrate t-value statistics. All activations thresholded at a voxelwise p<.05 FDR, k>5 voxels, masked by IFG and OFC regions of interest (k=9635 voxels in total). IFG: inferior frontal gyrus; tri: *pars triangularis*; op: *pars opercularis*; orb: *pars orbitalis*; OFC: orbitofrontal cortex; med: medial; mid: middle; sup: superior.

**Supplementary Table 1: Results for each factor of the modelling of accuracy data using generalized linear mixed-effects.**

| **Effect** | **Chi-square value** | **Degrees of freedom** | ***p*-value** |
|---|---|---|---|
| Species | 222.78 | 3 | <.001*** |
| Affective context | 4.64 | 2 | .099. |
| Species*Affective context | 12.78 | 6 | .047* |
| Mean of F0 | 0.14 | 1 | .712 |
| Std of F0 | 2.94 | 1 | .087. |
| Mean of Energy | 0.00 | 1 | .995 |
| Std of Energy | 0.01 | 1 | .937 |

F0: voice fundamental frequency; Energy: voice spectral energy; . $p<.10$, *$p<.05$, ***$p<.001$.

**Supplementary Table 2: Results for each factor and each species of the modelling of confusion data using linear mixed-effects.**

| Effect | Chi-square value | Degrees of freedom | *p*-value |
|---|---|---|---|
| **Human confusion** | | | |
| Species | 1.65 | 3 | .648 |
| Affective context | 0.21 | 2 | .090. |
| Species*Affective context | 4.65 | 6 | .589 |
| Mean of F0 | 0.03 | 1 | .870 |
| Std of F0 | 0.72 | 1 | .398 |
| Mean of Energy | 0.07 | 1 | .788 |
| Std of Energy | 1.18 | 1 | .277 |
| **Chimpanzee confusion** | | | |
| Species | 265.79 | 3 | <.001*** |
| Affective context | 6.44 | 2 | <.039* |
| Species*Affective context | 8.38 | 6 | .211 |
| Mean of F0 | 0.02 | 1 | .887 |
| Std of F0 | 0.10 | 1 | .748 |
| Mean of Energy | 0.37 | 1 | .542 |
| Std of Energy | 0.34 | 1 | .561 |
| **Bonobo confusion** | | | |
| Species | 107.96 | 3 | <.001*** |
| Affective context | 7.62 | 2 | .022* |
| Species*Affective context | 11.00 | 6 | .088. |
| Mean of F0 | 4.56 | 1 | .033* |
| Std of F0 | 6.65 | 1 | .009** |
| Mean of Energy | 0.02 | 1 | .874 |
| Std of Energy | 0.09 | 1 | .763 |
| **Macaque confusion** | | | |
| Species | 469.68 | 3 | <.001*** |
| Affective context | 2.25 | 2 | .325 |
| Species*Affective context | 6.27 | 6 | .393 |
| Mean of F0 | 4.96 | 1 | .025* |
| Std of F0 | 4.38 | 1 | .036* |
| Mean of Energy | 1.15 | 1 | .284 |
| Std of Energy | 0.95 | 1 | .329 |

F0: voice fundamental frequency; Energy: voice spectral energy; . $p<.10$, *$p<.05$, **$p<.01$, ***$p<.001$.

**Supplementary Table 3: Peak MNI coordinates of the correlates and anti-correlates of the probability of correctly classifying each species (voxel-wise *p*<.05 FDR, k>5 voxels, masked by bilateral IFG and OFC)**

| **Correlates** | | | | | | |
|---|---|---|---|---|---|---|
| **Region** | **Hemisphere** | **MNI x** | **MNI y** | **MNI z** | **T value** | **Voxels** |
| $OFC_{med}$ | L | -8 | 46 | -6 | 4.15 | 227 |
| $IFG_{tri}$ | R | 54 | 36 | -2 | 3.36 | 16 |
| $OFC_{med}$ | L | -30 | 16 | -22 | 3.33 | 6 |
| $IFG_{op}$ | R | 60 | 16 | 10 | 3.12 | 13 |
| **Anti-correlates** | | | | | | |
| **Region** | **Hemisphere** | **MNI x** | **MNI y** | **MNI z** | **T value** | **Voxels** |
| $IFG_{tri}$ | L | -46 | 38 | 8 | 8.15 | 1919 |
| $IFG_{tri}$ | R | 36 | 28 | 12 | 7.22 | 403 |
| $IFG_{op}$ | R | 40 | 4 | 26 | 4.56 | 53 |
| $IFG_{orb}$ | R | 26 | 34 | -8 | 4.54 | 53 |
| $IFG_{orb}$ | R | 24 | 42 | -12 | 3.41 | 5 |
| $OFC_{sup}$ | L | -40 | 54 | -2 | 3.18 | 5 |

IFG: inferior frontal gyrus; OFC: orbitofrontal cortex; tri: *pars triangularis*; med: medial; op: *pars opercularis*; orb: *pars orbitalis*; sup: superior.

**Supplementary Table 4: Peak MNI coordinates of conjunction analyses for non-human primate vocalization processing and correct categorization as compared to human voice (voxel-wise *p*<.05 FDR, k>5 voxels, masked by bilateral IFG and OFC)**

| **Conjunction: [Chimpanzee > Human] > [Bonobo > Human] > [Macaque > Human] (processing)** | | | | | | |
|---|---|---|---|---|---|---|
| **Region** | **Hemisphere** | **MNI x** | **MNI y** | **MNI z** | **T value** | **Voxels** |
| $IFG_{op}$ | L | -46 | 4 | 28 | 6.62 | 1337 |
| $IFG_{tri}$ | R | 40 | 32 | 14 | 5.53 | 139 |
| $IFG_{op}$ | R | 44 | 4 | 26 | 4.29 | 29 |
| $IFG_{orb}$ | L | -28 | 34 | -12 | 4.17 | 22 |
| $IFG_{tri}$ | R | 48 | 34 | 24 | 3.50 | 7 |
| **Conjunction: [Chimpanzee > Human] > [Bonobo > Human] > [Macaque > Human] (correct responses)** | | | | | | |
| **Region** | **Hemisphere** | **MNI x** | **MNI y** | **MNI z** | **T value** | **Voxels** |
| $IFG_{tri}$ | L | -36 | 28 | 16 | 4.76 | 89 |

IFG: inferior frontal gyrus; tri: *pars triangularis*; op: *pars opercularis*; orb: *pars orbitalis*.
